## Supplementary Figure for "On the identification of differentially-active transcription factors from ATAC-seq data"

Supplementary Figures

Gerbaldo, Sonder et al.

31 July, 2024

### Supplementary Figure 1

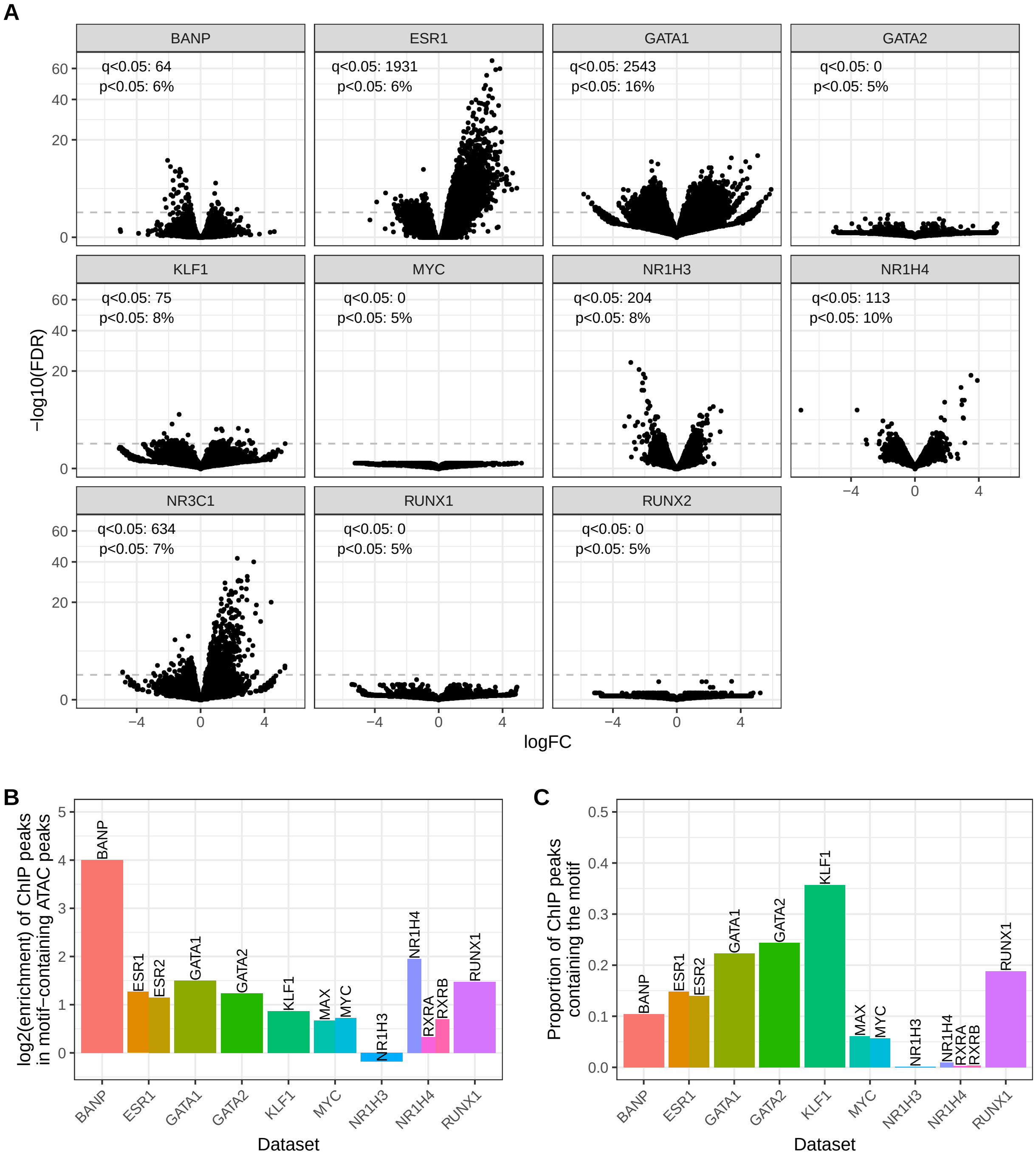


**Supplementary Figure 1: Overview of the benchmark datasets and TFs.**
**A:** Volcano plot of the peak-level differential accessibility analyses, illustrating the extent and significance of changes upon treatment in each dataset. The dashed line represents a 0.05 FDR threshold, and indicated are the number of ATAC peaks passing this threshold, as well as the proportion of peaks with an uncorrected p-value lower than 0.05. **B:** Enrichment for experimental binding sites of the factor (main TF after which the datasets are named) in ATAC peaks containing matches for the respective motifs, versus all ATAC peaks. **C:** Proportion of ChIPseq peaks (overlapping ATAC peaks) that contain matches for the respective motif.

#### Supplementary Figure 2

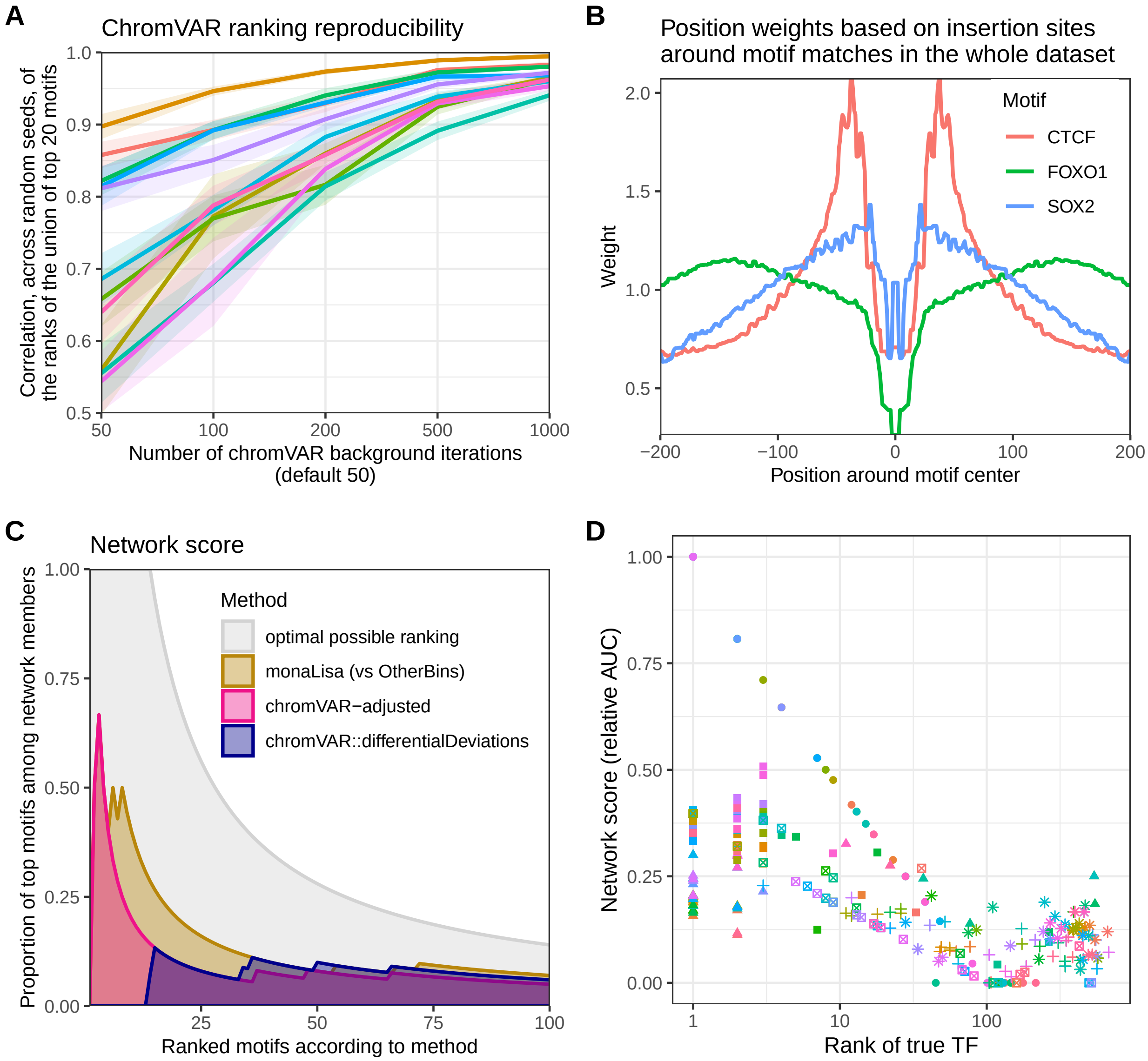


**Supplementary Figure 2 A: Reproducibility of chromVAR-based differential accessibility analysis.** Each color represents a dataset. The line and shaded area respectively represent the mean and standard error of the pairwise correlations across 7 independent runs of the chromVAR(z-score)>limma pipeline for each dataset and setting, of the ranks of the (union of) top 20 motifs. **B: Illustration of the motif-specific position weights for the insertionModel method. C: Illustration of the computation of the network score.** The motifs are ranked for each method, and for each k=1:100 the proportion of top k motifs that are part of the known network of the true TF is computed. The area under the curve is then reported, relative to the AUC of the best possible motif ordering for that factor (gray area). The example plotted here is from the GATA1 dataset. **D: Agreement between the two rank-based metrics.** Each point represents the results of a method (colour) in a dataset (shape). The network score is not entirely comparable across datasets. Beside this effect, and except at very low (1-3) or high (>100) ranks (i.e. where the ranks stop being discriminatory), there is a good agreement between the two metrics. Capping ranks to 100, the overall correlation between between ranks and network score is -0.61, and the median correlation across datasets is -0.75.

### Supplementary Figure 3

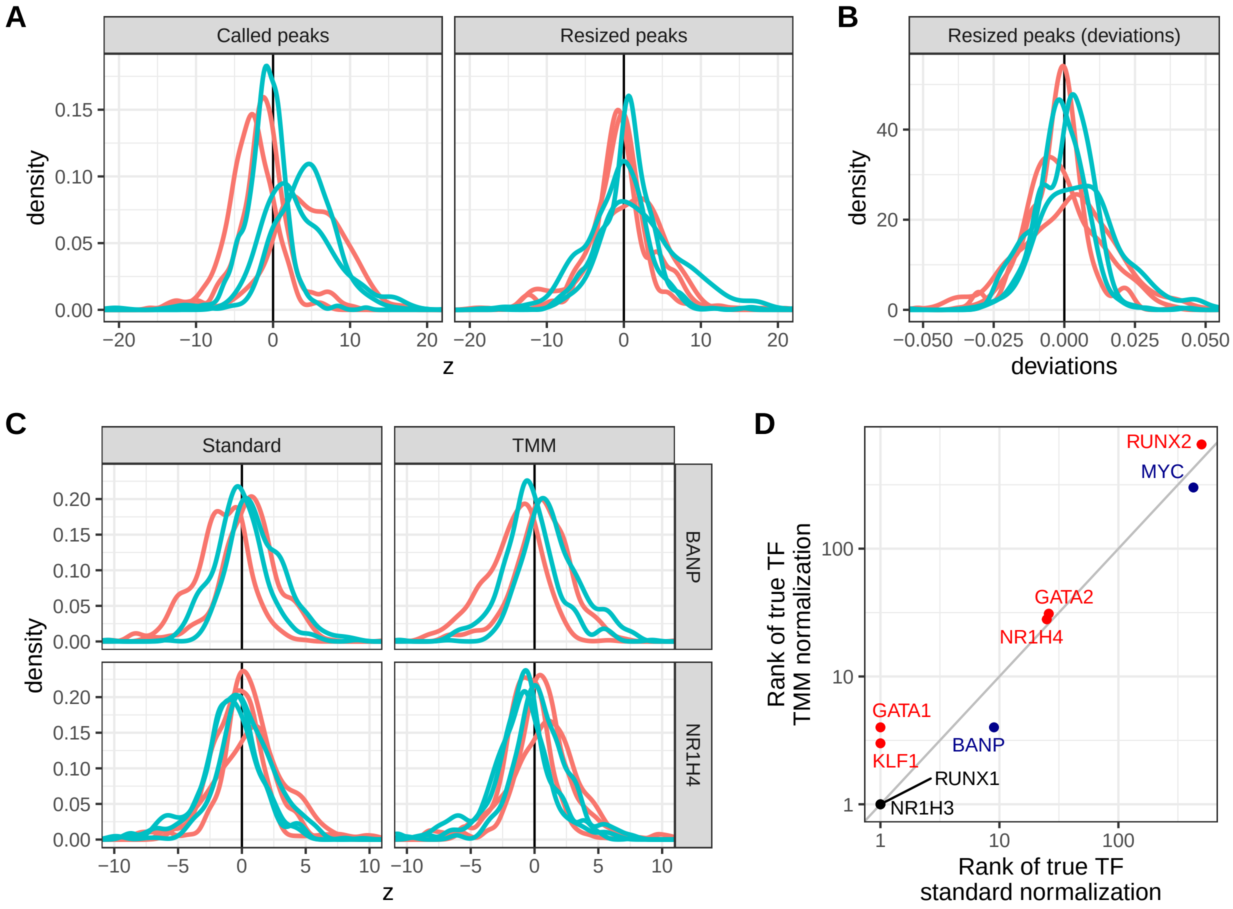


**Supplementary Figure 3: Differences in distributions of chromVAR z-scores across samples. A-B:** Example of a subset of ATAC samples from Caradonna, Paul and Marrocco (2022) (GSE200670), where different samples show different shapes of z-scores distributions across motifs/TFs. The colors indicate different experimental conditions. When peaks are not resized to all have the same width, as recommended, the distributions are often globally shifted (**A**, left). Even when peaks are resized (**A**, right), differences in the width of the z-scores distributions which are not related to experimental groups can persist in some datasets. This is not specific to the z-scores, but also present in the bias-corrected deviations (**B**). **C-D:** Effect on replacing the internal library size normalization performed by chromVAR with a TMM normalization. TMM normalization does not reduce the differences in z-score distributions (**C**), nor does it improve the rank of the true TF (**D**).

#### Supplementary Figure 4

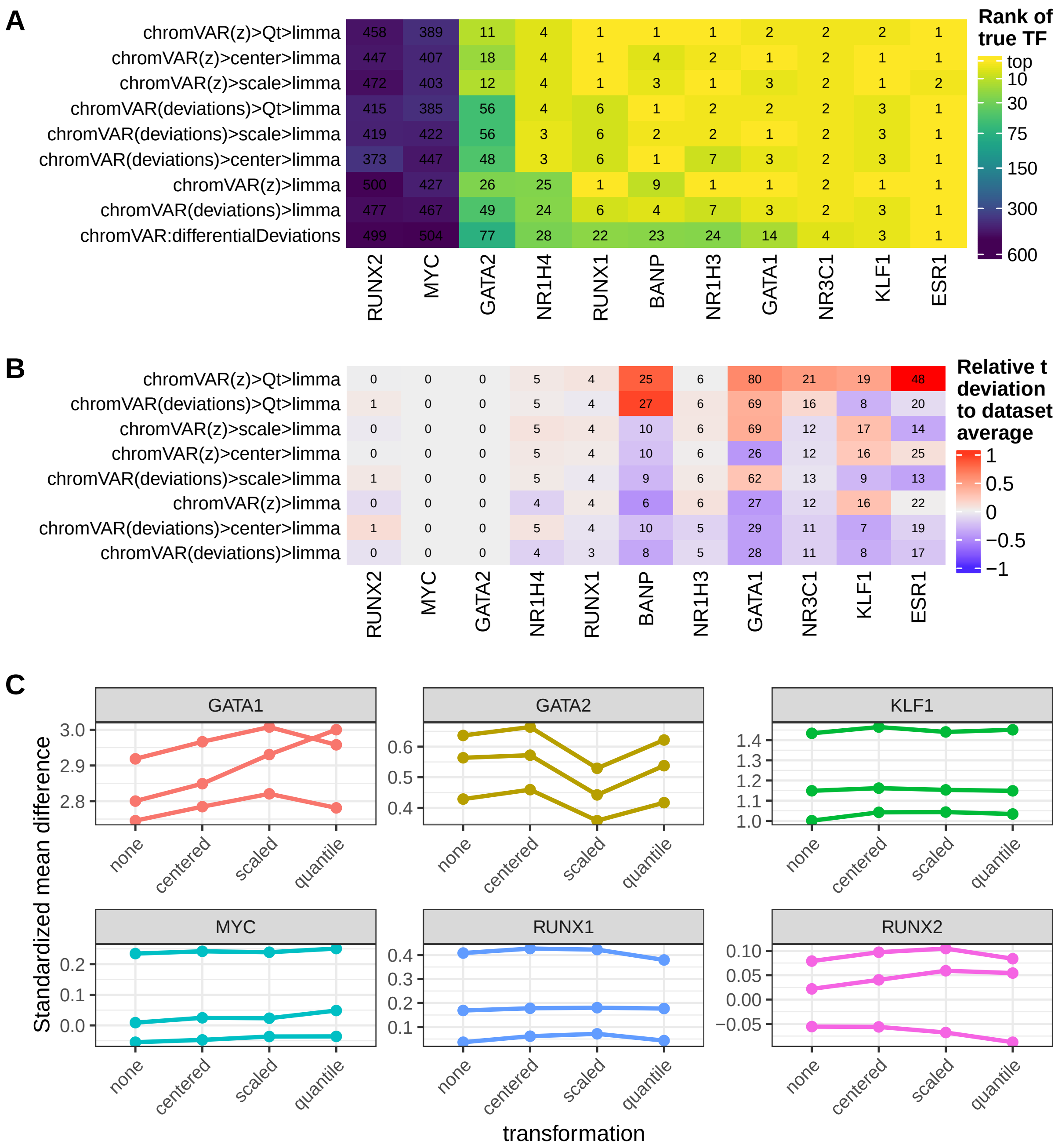


**Supplementary Figure 4: Impact of normalization of the chromVAR z-score distributions. A-B:** Effect on the rank of the true TF (**A**) and its *t*-value (**B**) of normalizing the chromVAR activity scores (i.e. z-scores or deviations) distributions across samples before running limma. In **B**, for datasets where a downregulation was expected, the *t*-values were inverted. The heatmap colors show the methods deviation relative to the dataset average, while the numbers indicate the actual (rounded) *t*-value. **C:** Impact on the single-cell z-scores. As a proxy to signal-to-noise ratio, we computed the standardized mean difference of the respective motif for each guide RNA (individual lines) to the mean of the control guide RNAs. Centering is always beneficial (Wilcoxon p-value<8e-06), unit-variance scaling is often beneficial (Wilcoxon p-value~0.07) but sometimes detrimental, and quantile normalization also tends to be beneficial (Wilcoxon p-value~0.04).

### Supplementary Figure 5

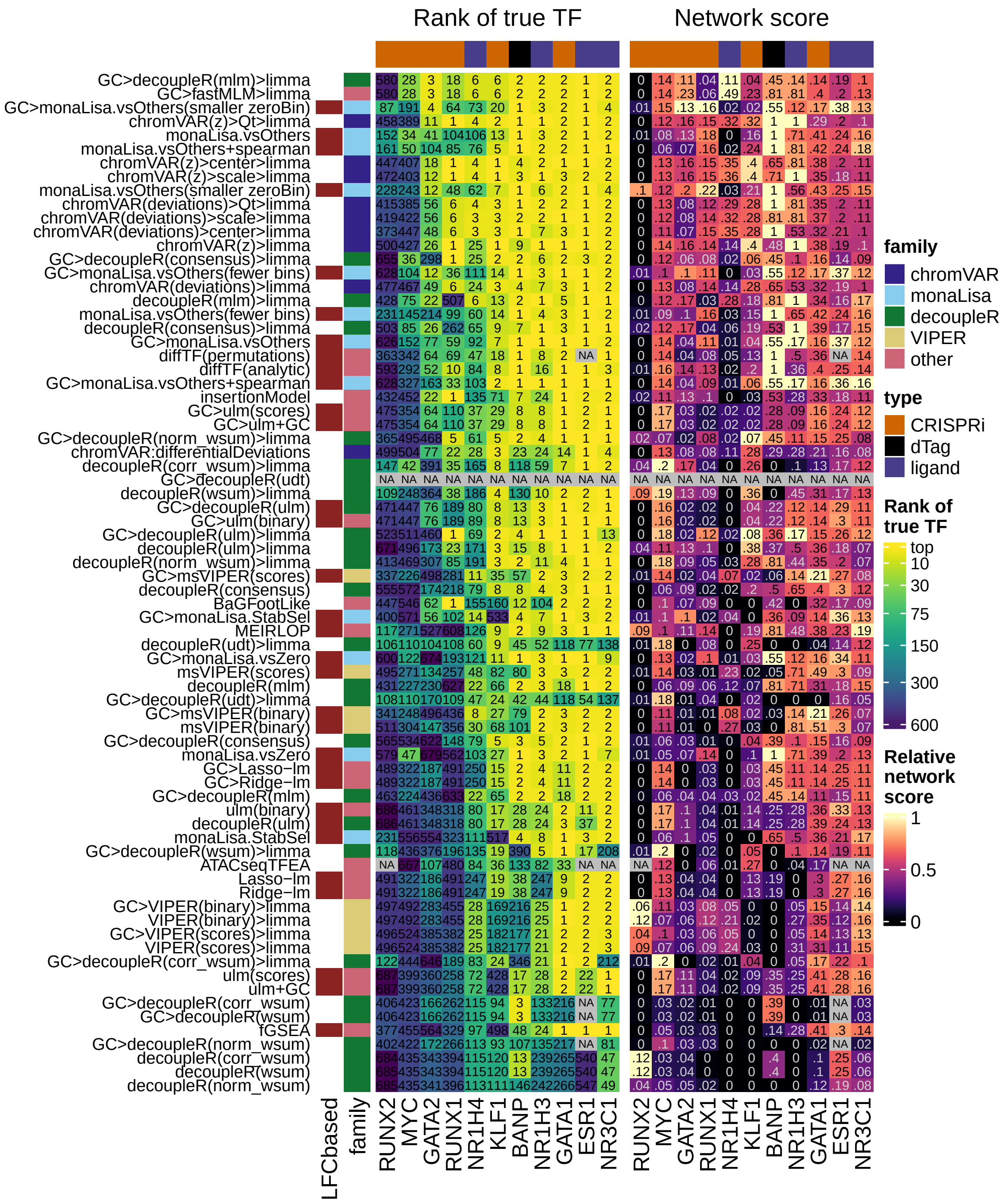


**Supplementary Figure 5: Rank-based metrics of all variants, including smooth quantile normalization in GC bins.** Same as Figure 1, but showing all method variants tested.

### Supplementary Figure 6

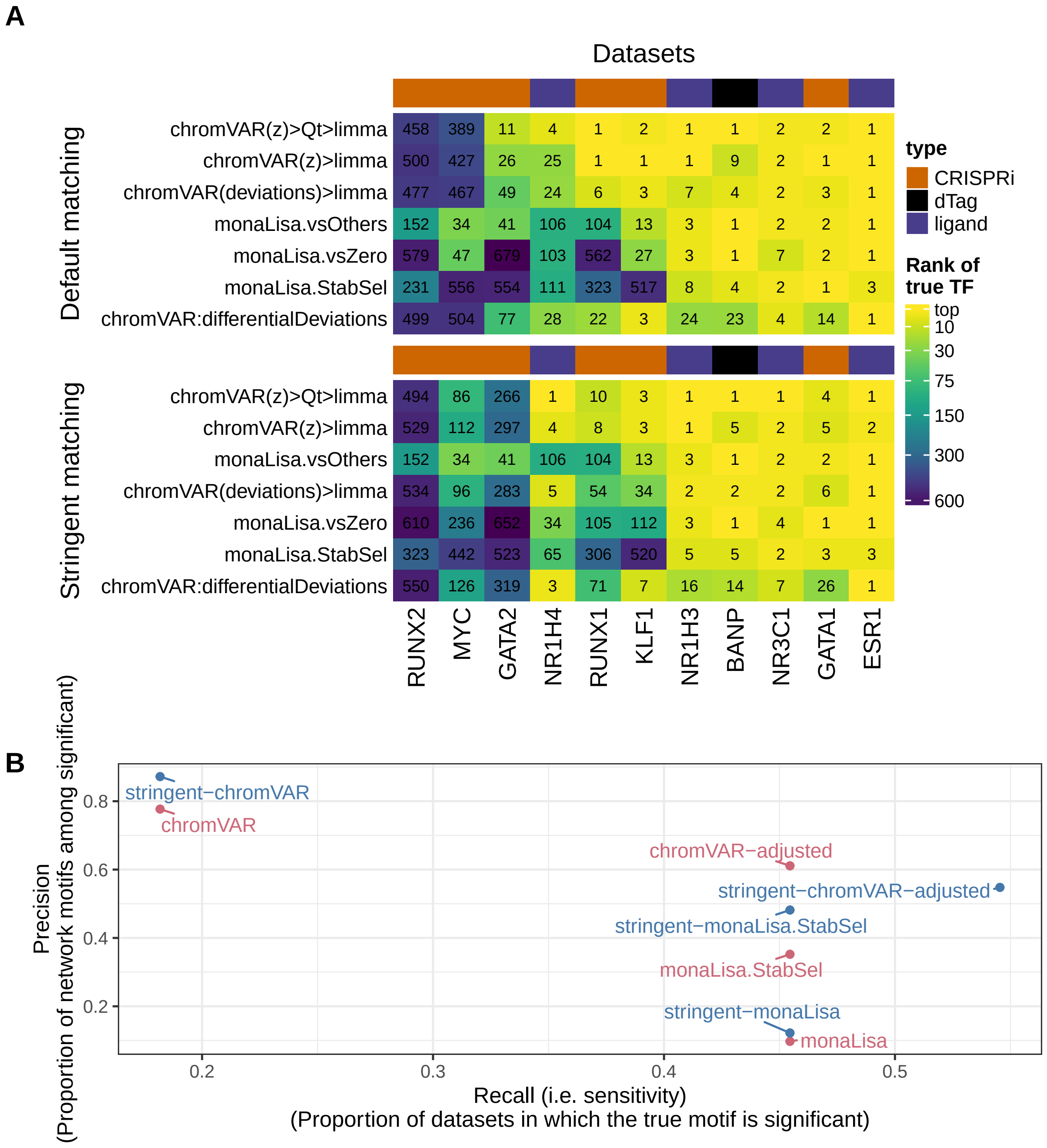


**Supplementary Figure 6: Impact of stringency in motif matching.** Rank of the true TF (**A**) and Precision-Recall (**B**) of selected methods, when using a stringent motif matching vs the default.

### Supplementary Figure 7

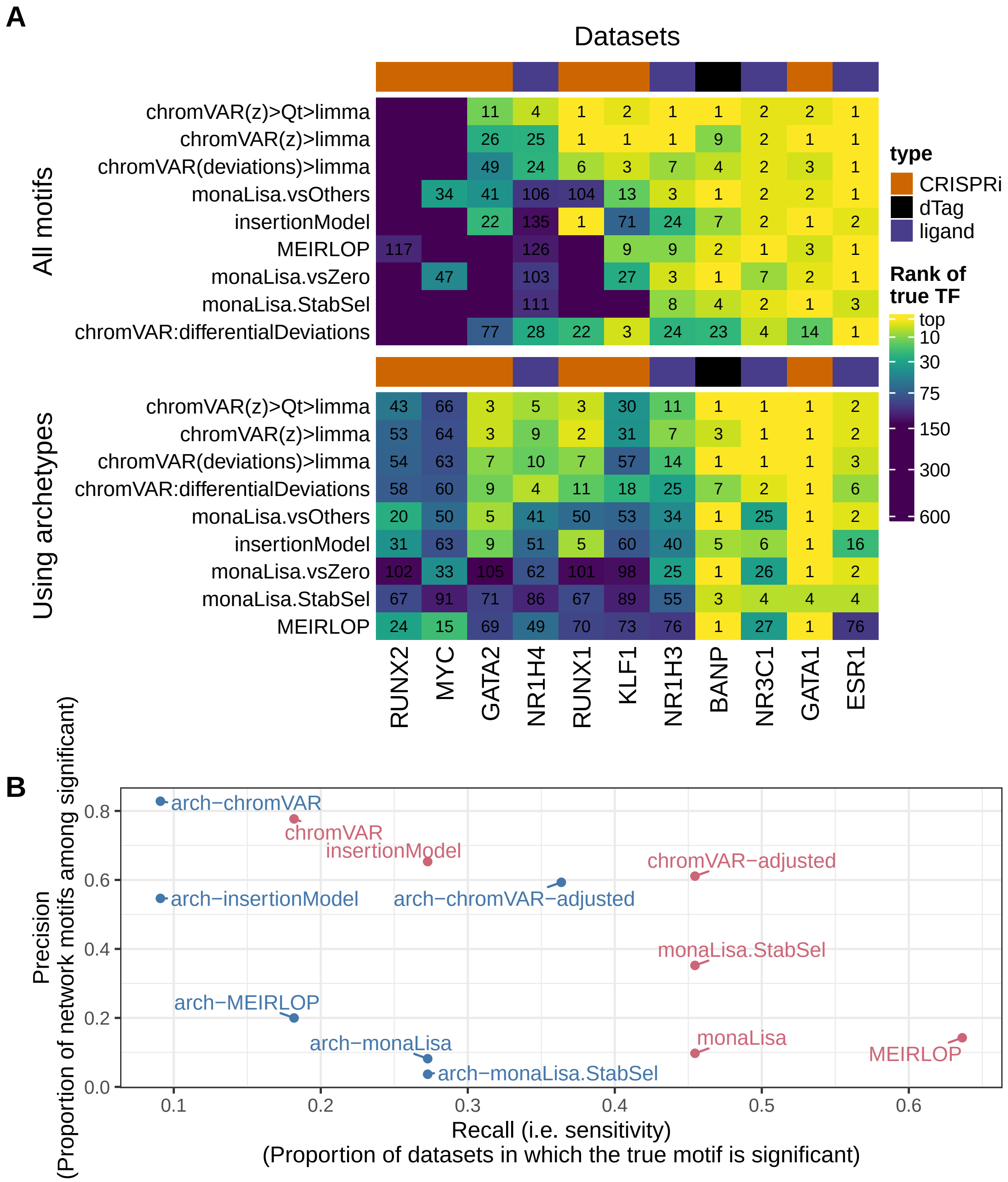


**Supplementary Figure 7: Impact of using motif archetypes.** Rank of the true motif/archetype (**A**) and Precision-Recall (**B**) of selected methods, when using motifs archetypes instead of the full collection of motifs. Given the difference in the size of the two sets of motifs, ranks were capped at 144 for the purpose of creating the heatmap.

### Supplementary Figure 8

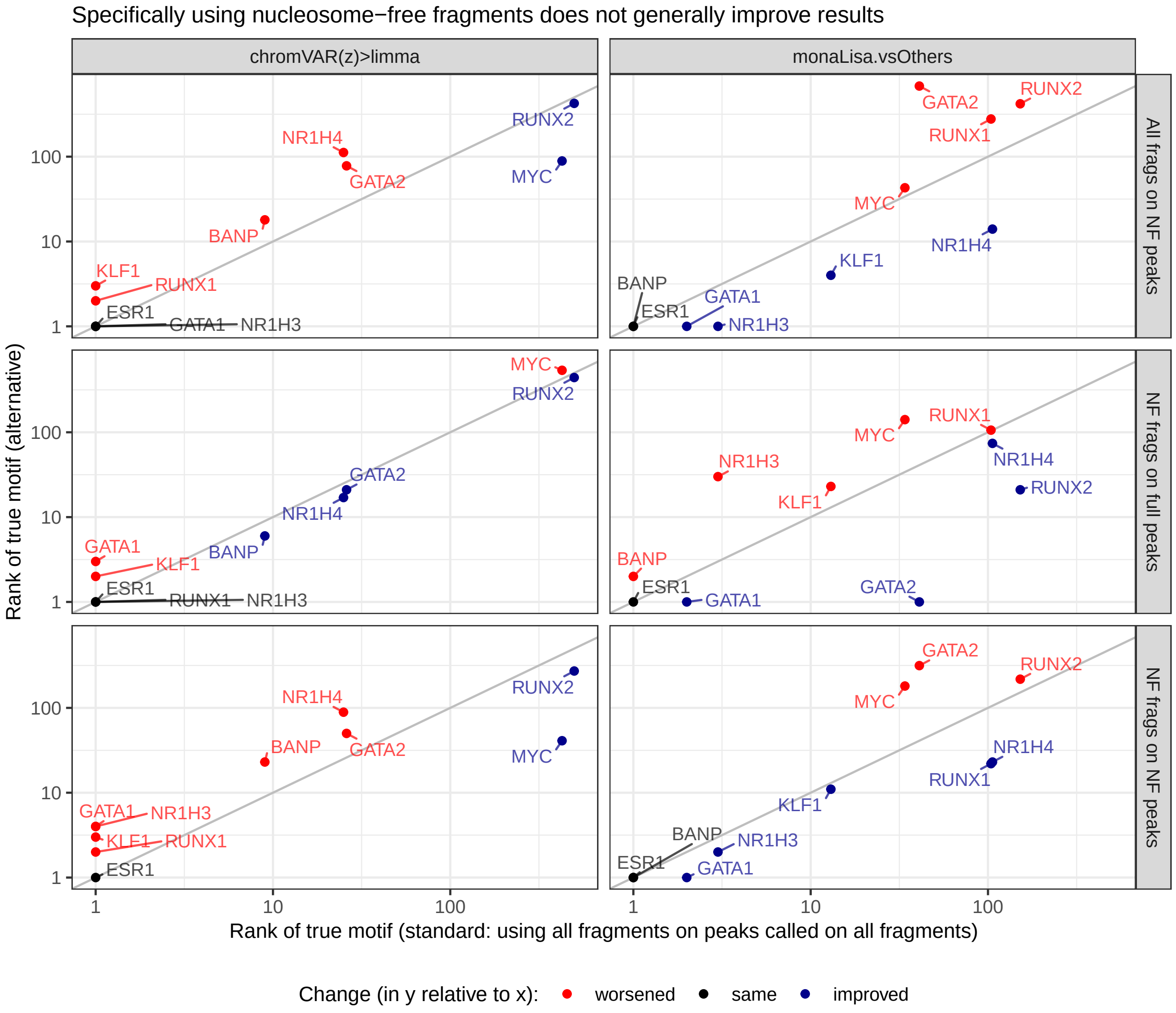


**Supplementary Figure 8: Impact of using nucleosome-free fragments.** Comparison of the rank of the true motif obtained by the top method of each family of approaches, when using alternative peaks or overlap counts based on nucleosome-free fragments. 'All frags on NF peaks' stands for counting overlaps with fragments of any size, but on peaks called on nucleosome-free (NF) fragments only. 'NF frags on NF peaks' stands for using only NF-fragments, i.e. counting overlaps of NF fragments on peaks called only on NF fragments. Finally, 'NF frags on full peaks' stands for counting overlaps of only NF fragments, but on the standard peaks (called using all fragments).

### Supplementary Figure 9

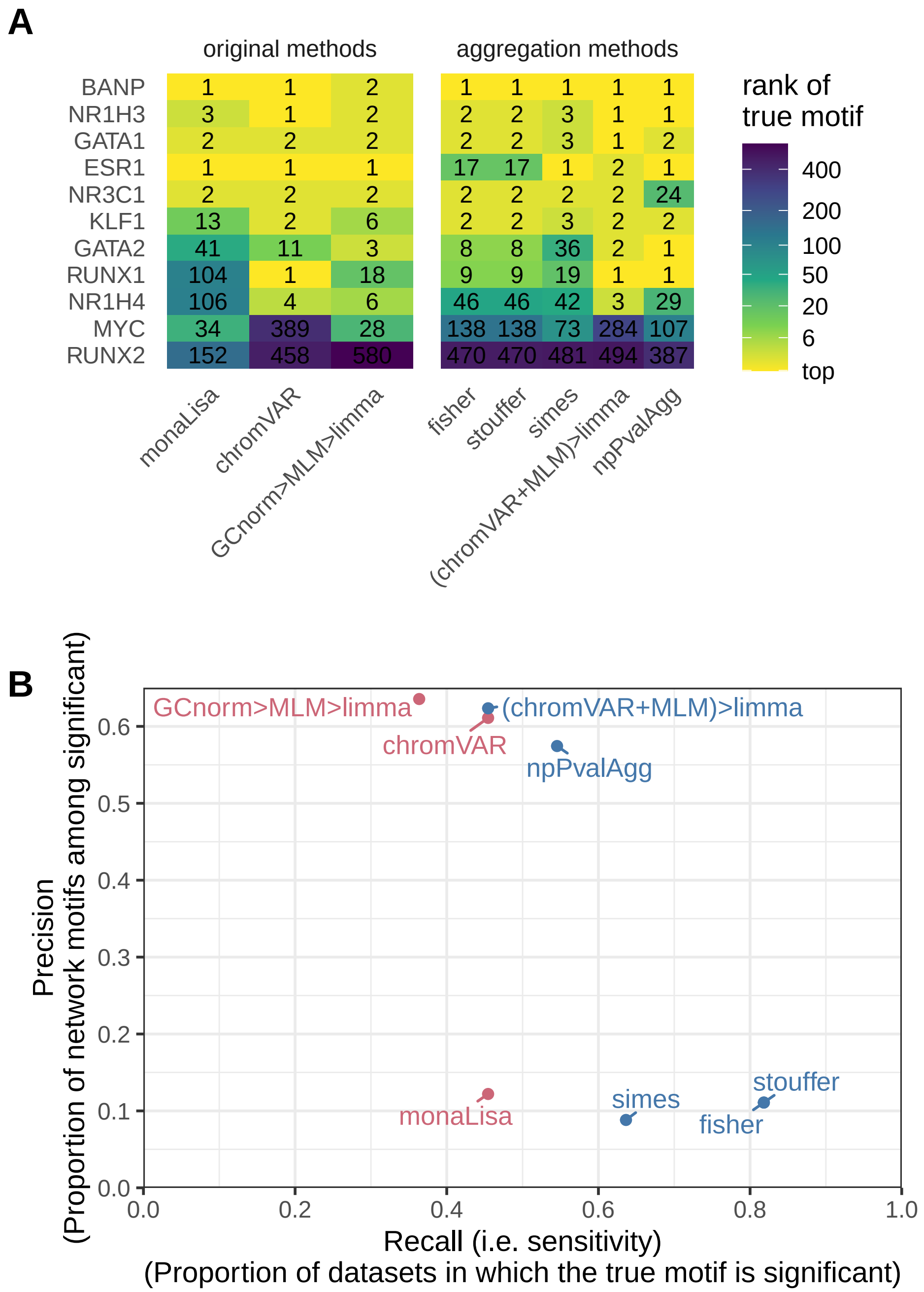


**Supplementary Figure 9: Simple aggregation of top methods' results does not improve inferences.** **A**. The results of the top method from each of the two families of approach (on the left) are compared to those of simple aggregation methods (on the right). Beside the established Fisher's, Stouffer's and Simes' methods, a rank-based permutation approach was tested, establishing the probability of having a sum of ranks across methods lower or equal to the observed one. We also tested using limma on merged per-sample activity scores from chromVAR and GCsmooth+MLM. **B**. Same comparison, in terms of Precision and Recall at adjusted p-value <= 0.05.

### Supplementary Figure 10

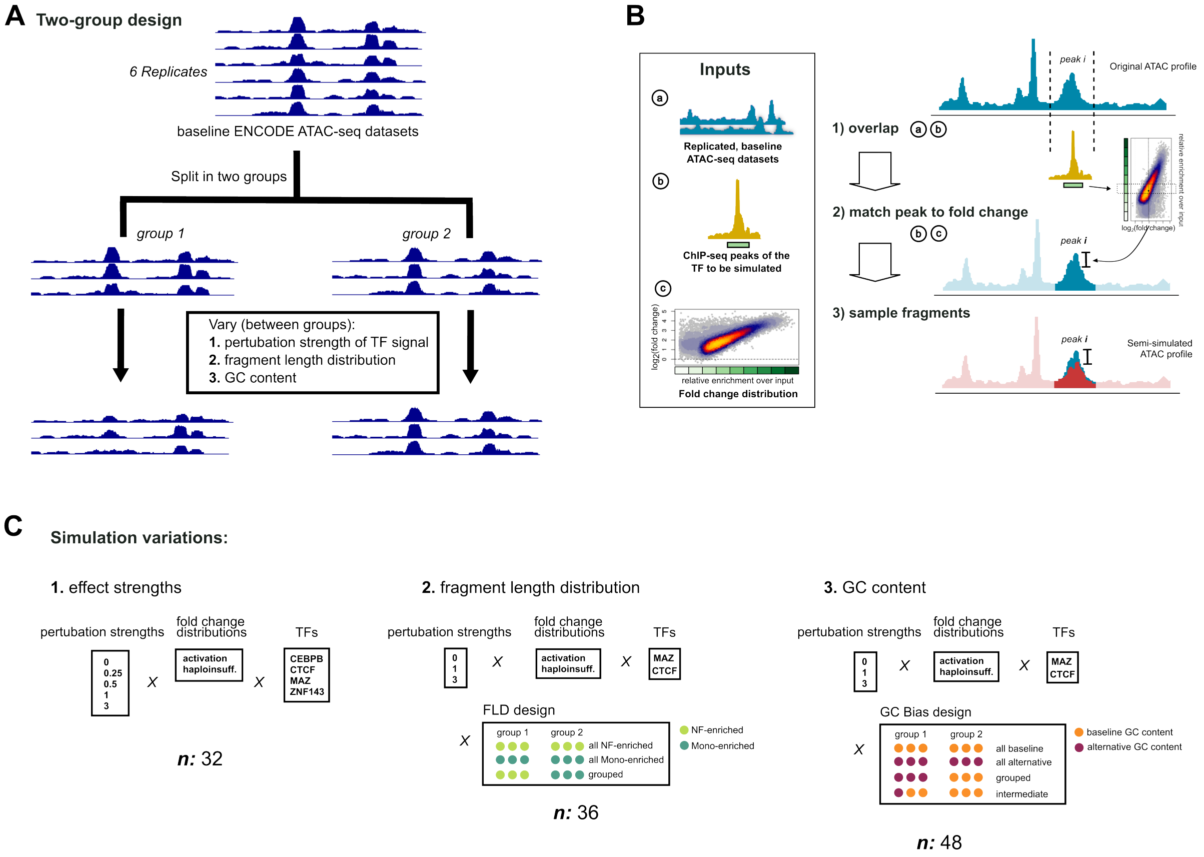


**Supplementary Figure 10: Overview of the semi-simulations.** **A:** 6 baseline ATAC-seq lymphoblastoid cell lines (LCLs) are divided into two groups. Differences will be introduced through downsampling of fragments in one of the groups, and eventual per-sample GC and fragment length biases are introduced. **B:** The same as show in main Fig 4A. The extent of downsampling is based on the enrichment over the input of the ChIP-seq peak, following one of two differential activity scenarios: an activation and a haploinsufficiency (see Fig 4B). The magnitude of the effect is further scaled up and down (perturbation strength). **C.** Settings used for the semi-simulated datasets. The resulting datasets can be divided into three groups, a first set where only the strength of the perturbation was varied by a factor (perturbation strength), a second and third one were additionally differences in respectively fragment length distribution and GC were introduced across samples.

### Supplementary Figure 11

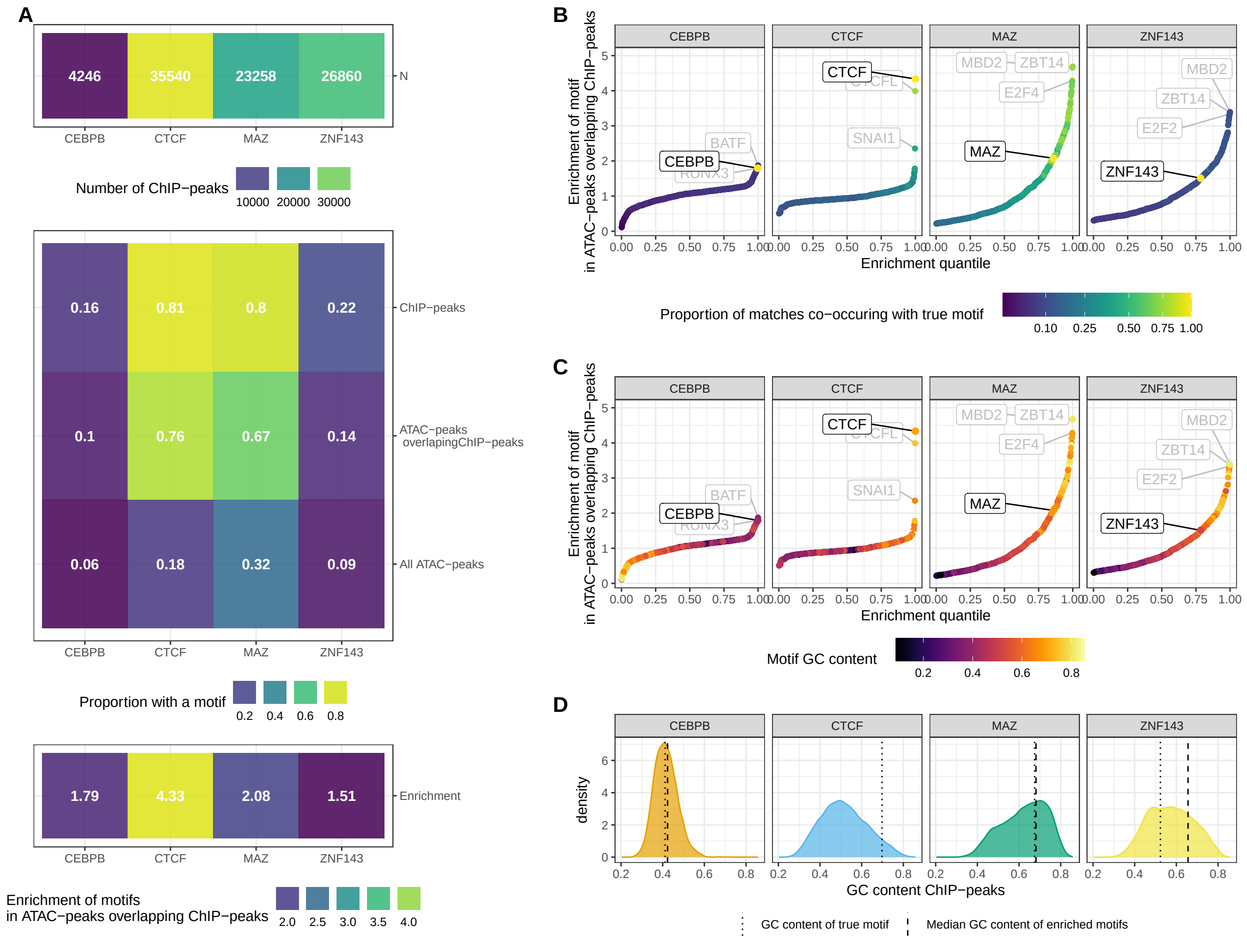


**Supplementary Figure 11: Overview of the simulated TFs.** Data shown correspond to the semi-simulated datasets with a pertubation strength of 1 in the activation paradigm. **A:** Number of peaks, proportion of ChIP-seq and/or ATAC-seq peaks with a motif, and fold-enrichment of ATAC-peaks overlapping ChIP-seq peaks (i.e. bound peaks) for the motif across the four TFs. **B-C:** Enrichment of all tested motifs among the ATAC-seq peaks overlapping ChIP-seq peaks (relative to all ATAC-seq peaks), with the true motif labeled in black. The dots are colored either by co-occurence with the true motif (**B**) or by GC content, i.e. the summed probabilities of G and C across positions in the position probability matrix divided by the total sum (**C**). **D:** GC content distributions of the ChIP-seq peaks of each TF. With dotted lines the GC content of the true motif is indicated, with dashed lines the median GC content of the motifs showing stronger enrichment in the ChIP-seq peaks than the true motif.

### Supplementary Figure 12

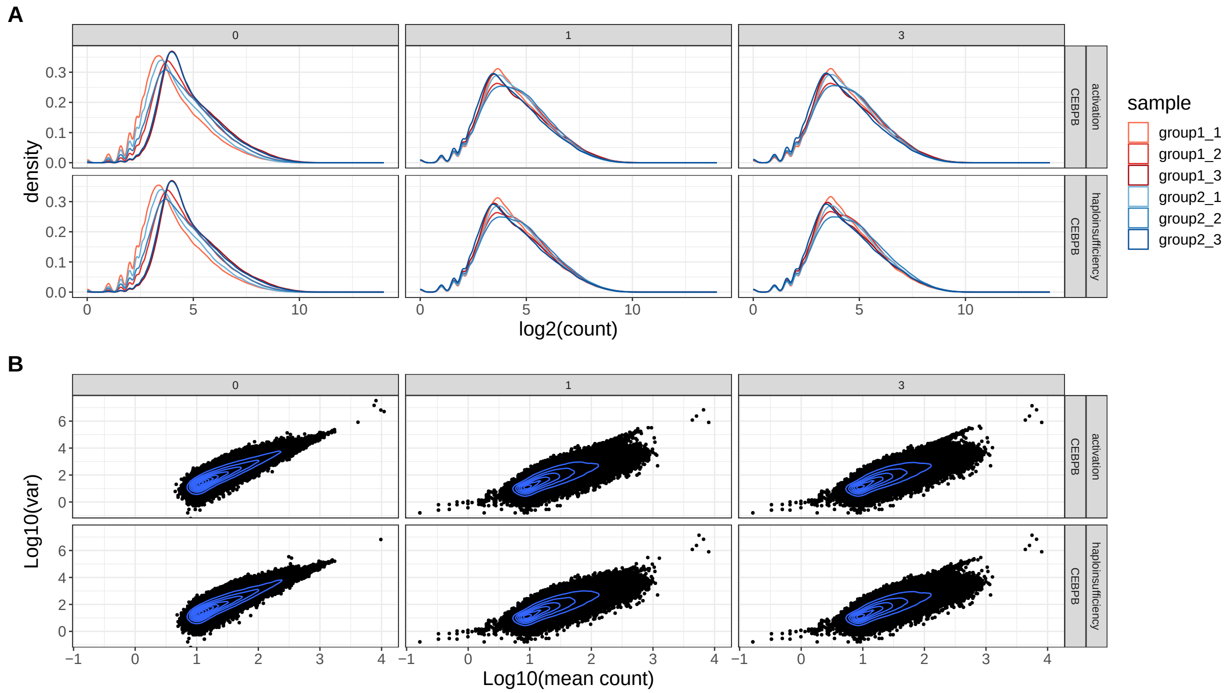


**Supplementary Figure 12: Distribution of fragment counts and variance of the semi-simulations.** **A:** Per peak fragment count distributions of the original (dataset: “0”) and simulated samples (datasets: “1”, “3”) with 1-fold and 3-fold perturbation strengths, for an example TF (CEBPB) across both paradigms. **B:** Mean-variance relationship of the original and simulated samples.

### Supplementary Figure 13

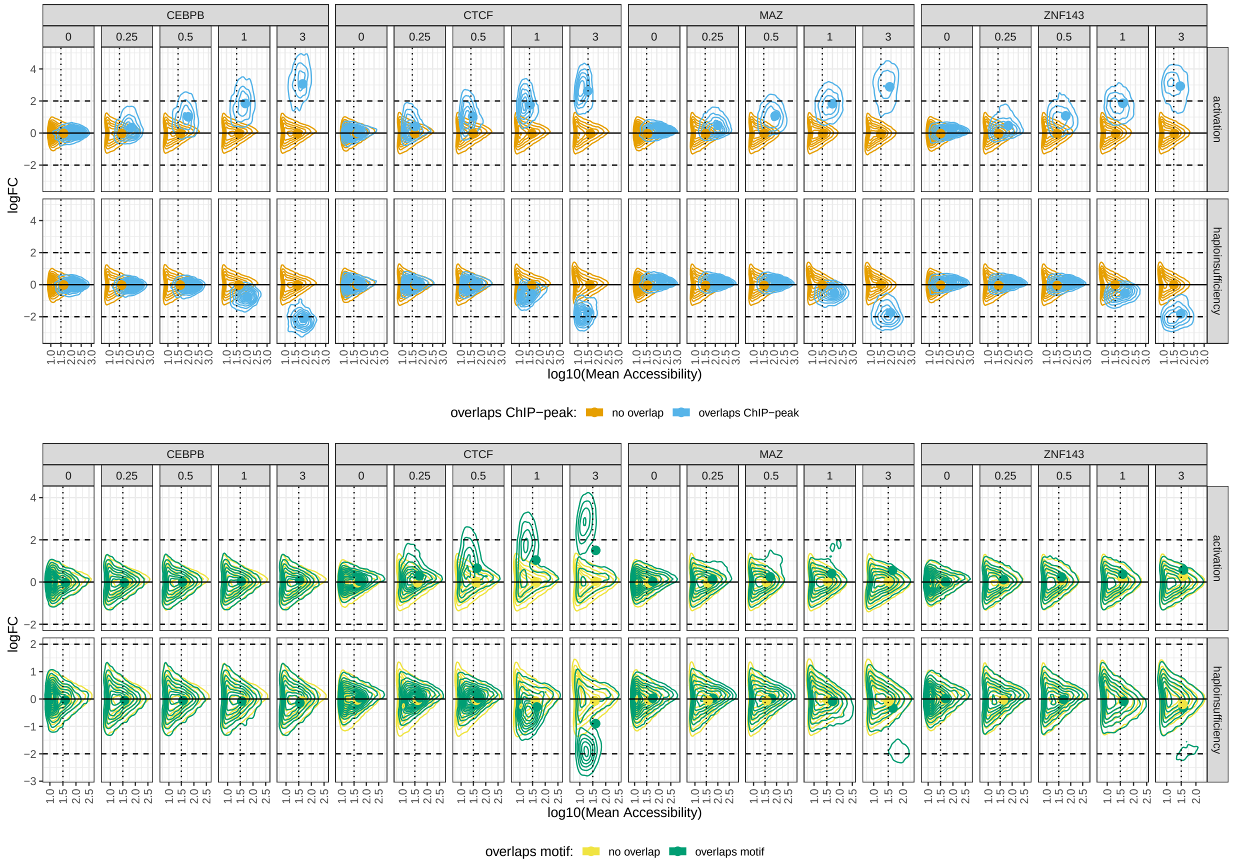


**Supplementary Figure 13: MA plots of the simulated effects.** For each factor, simulated paradigm and perturbation strength, the distribution of peaks log2FC of raw fragment counts between conditions as a function of mean accessibility is plotted as 2d kernel density, separating either (**A**) ATAC-seq peaks overlapping a ChIP-seq peak or not, or (**B**) ATAC-seq peaks containing a motif or not for the TF. For visibility, the peaks were subsampled to a more comparable size across the semi-simulated datasets (1.2$×$10ˆ5 peaks per dataset). Colored Points indicate the means of the respective distributions.

### Supplementary Figure 14

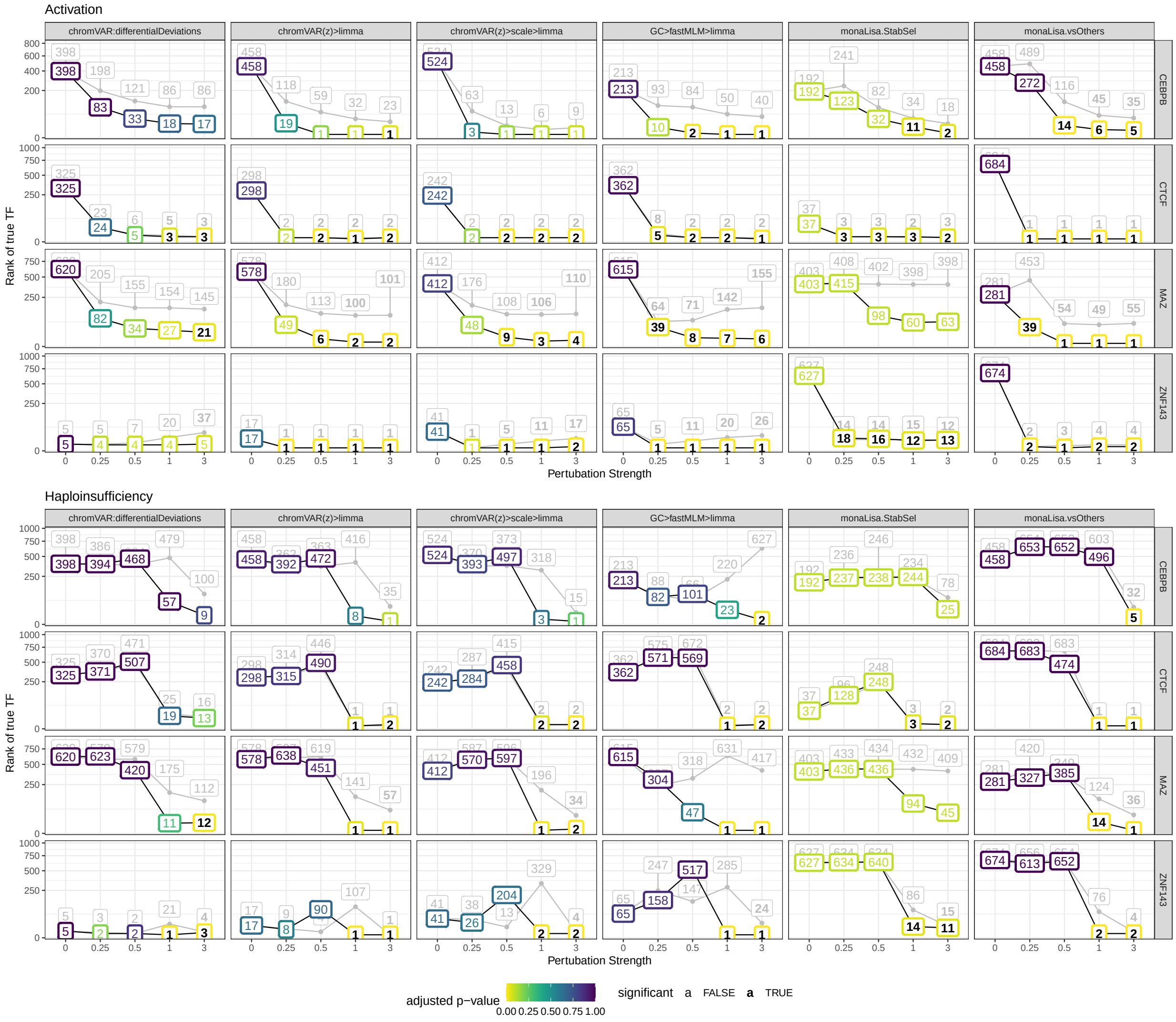


**Supplementary Figure 14: Positive control of the semi-simulations.** Shown are ranks and significance assigned by each method to the true motif across simulation parameters (TF, paradigm and perturbation strength). In light gray are the normal simulations (as shown in main Fig 4), while dark line and colored boxes correspond to the positive controls where a difference between the groups was only introduced in the peaks overlapping a motif of the true TF.

### Supplementary Figure 15

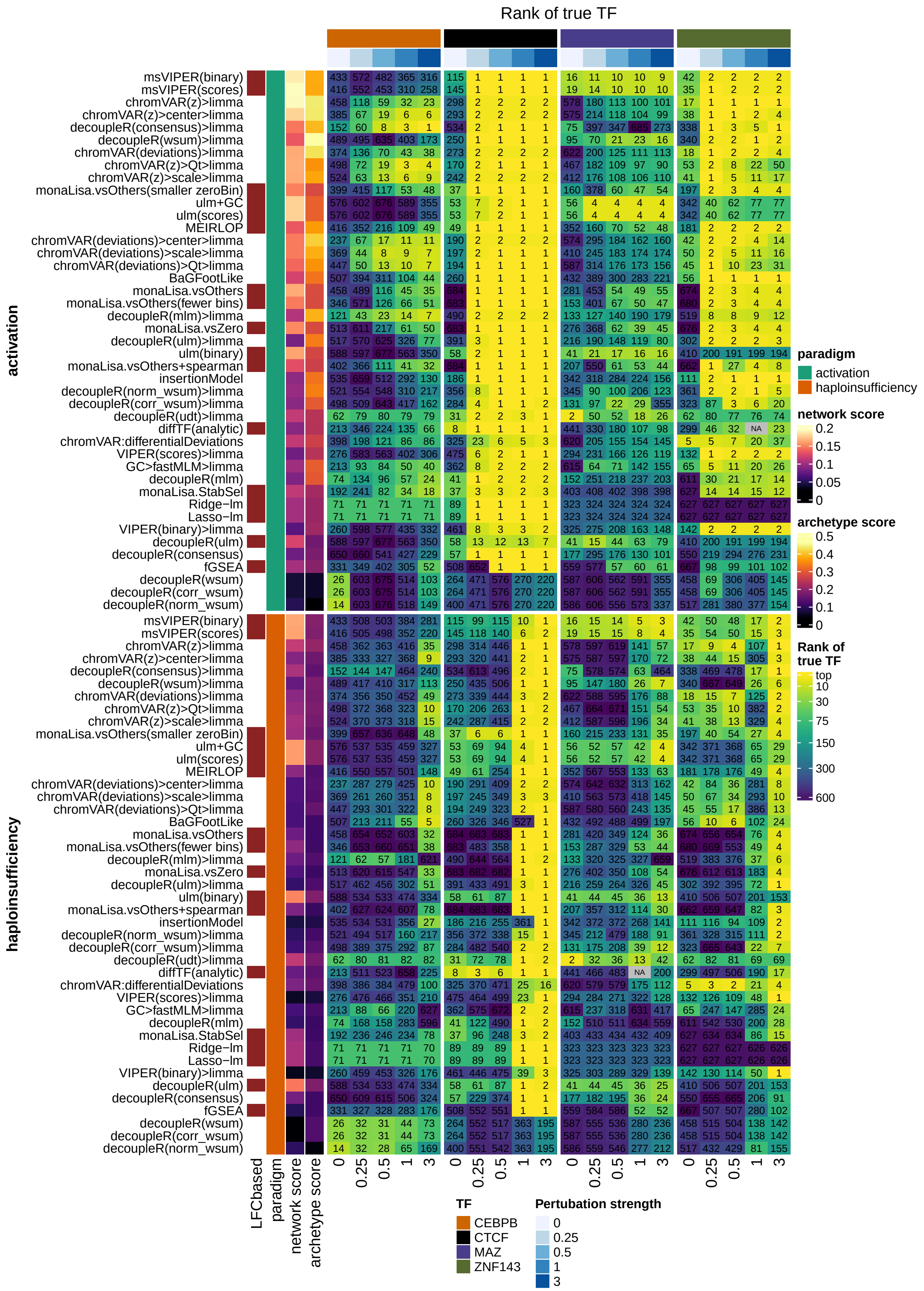

**Supplementary Figure 15: Heatmap of performance of methods on the semi-simulated datasets.** As in main Fig 1, the heatmap shows the ranks the methods achieved across different simulated pertubation strengths and fold change distributions (activation and haploinsufficiency). The annotation bar to the left shows the mean network and archetype scores on pertubation strengths greater than zero. Ranking of the methods was performed as in Fig 1 and described in the methods section using all the datasets with a pertubation strength greater than zero. Where needed imputation of missing values for the ranking was performed as previously described using only datasets with the same pertubation strength and fold change distribution. ATACseqTFEA was excluded for runtime reasons and diffTF could not be run on two of the datasets.

### Supplementary Figure 16

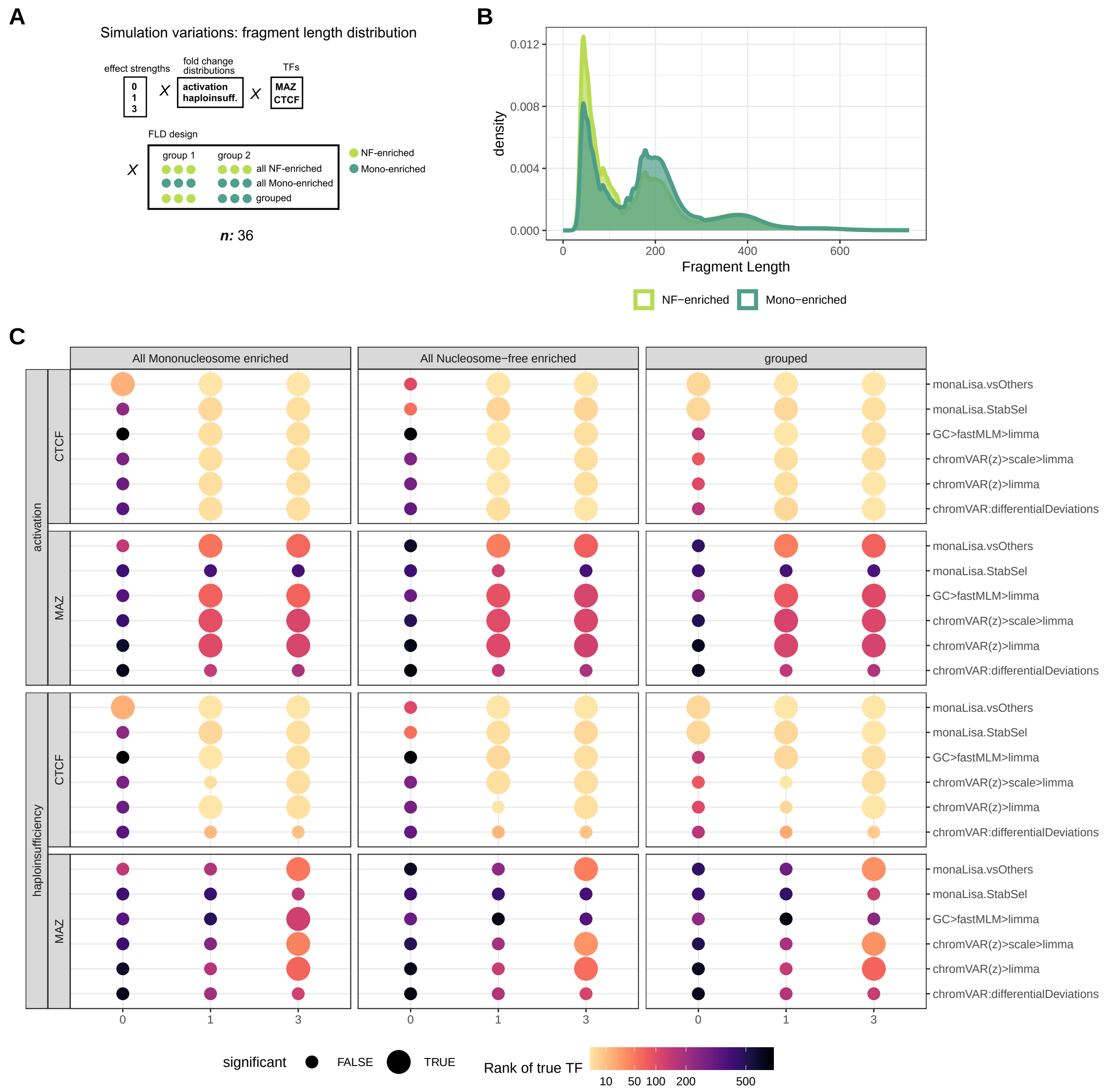


**Supplementary Figure 16: Robustness to simulated variations in fragment length.** In addition to the simulated TF effect, two different fragment length distributions (**B**) were simulated by differential sampling of the fragments according to their length for the different samples as in scheme **A**. The distribution in light green corresponds to sampling mononucleosome containing fragments with higher probability, whereas for the distribution in dark green nucleosome-free fragments were sampled preferentially. The results of the methods are shown in **C**. Colors of the points correspond to the rank of the true TF obtained, point size if the corresponding adjusted p-value was found to be significant (adjusted p-value <= 0.05).

### Supplementary Figure 17

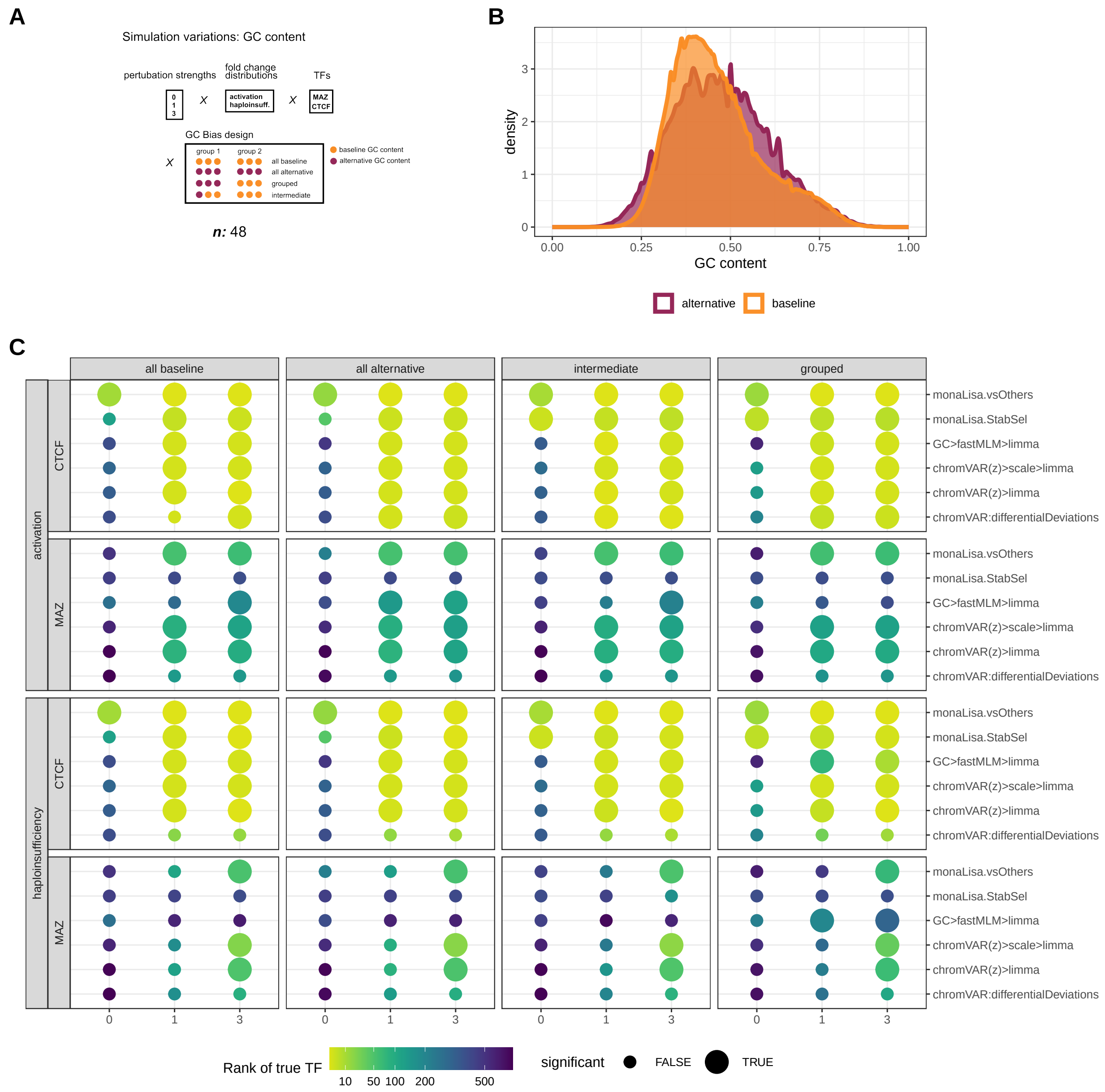


**Supplementary Figure 17: Robustness to simulated variations in GC content.** In addition to the simulated TF effect, two different GC content distributions (**B**) were simulated by differential sampling of the fragments according to their GC content for the different samples as in scheme **A**. The results of the methods are shown in **C**. Colors of the points correspond to the rank of the true TF obtained, point size if the corresponding adjusted p-value was found to be significant (adjusted p-value <= 0.05).

#### Supplementary Figure 18

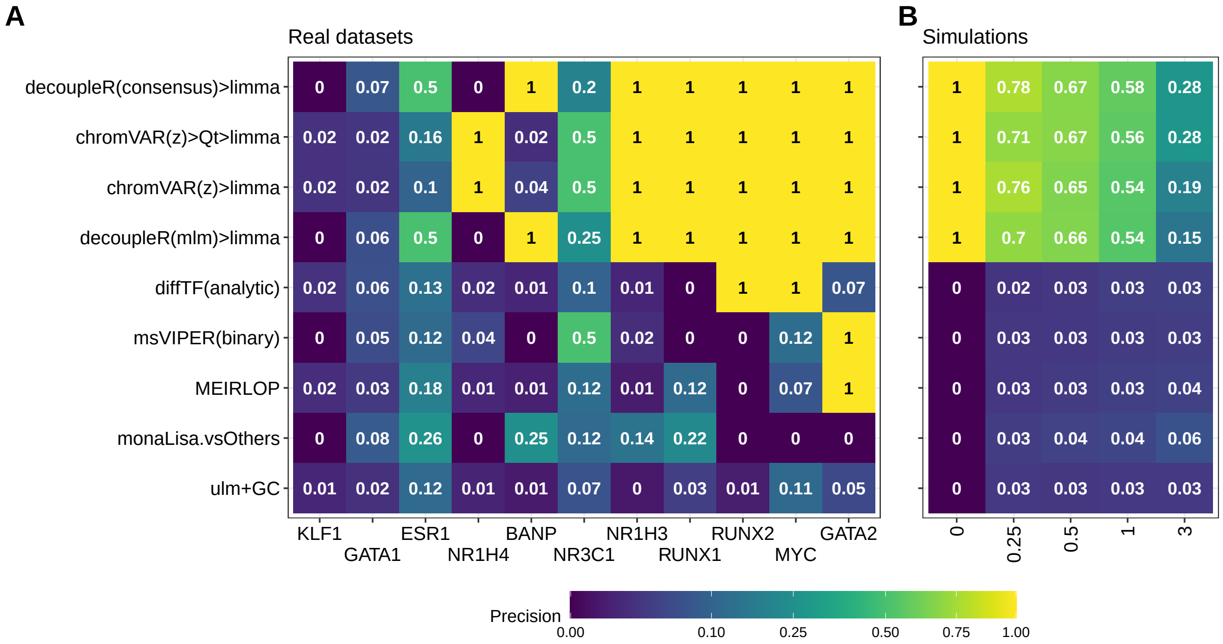


**Supplementary Figure 18: Estimated precisions per dataset.** Heatmap of precision estimates obtained per dataset and method on the 11 real datasets (**A**), as well as averaged across each perturbation strength of the semi-simulations (**B**).

#### Supplementary Figure 19

A

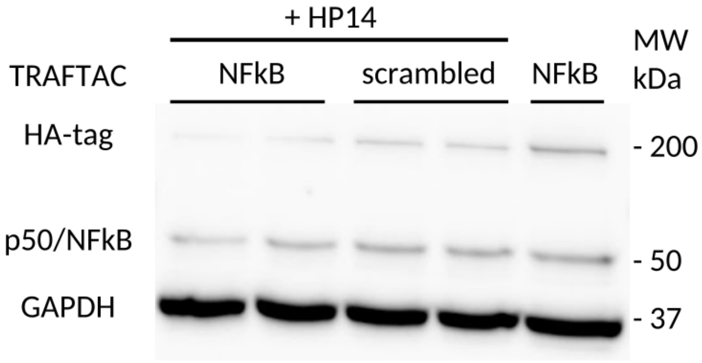


B

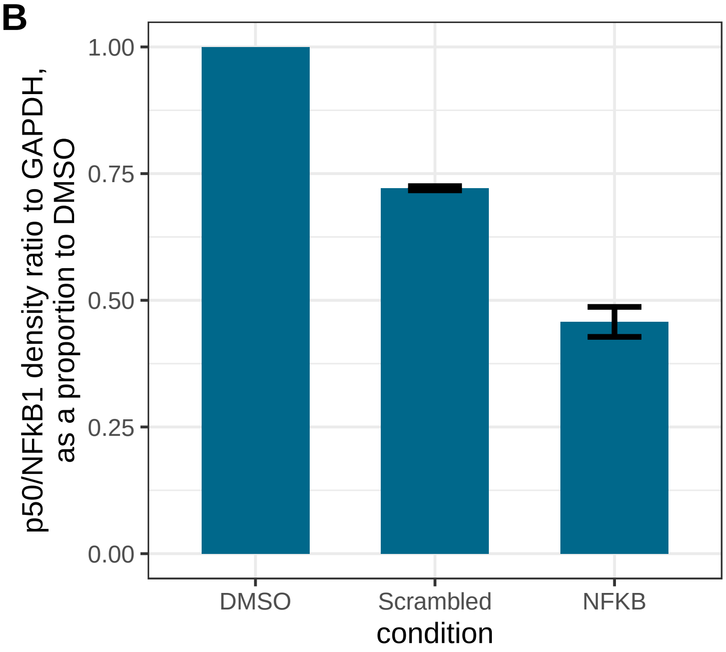


**Supplementary Figure 19: Successful downregulation of NFkB1 at the protein-level using the TRAFTAC.** Western Blot (**A**) and derived densitometry (**B**) of NFkB1 (aka p50) in HEK cells transfected with the dCas9-HT7 plasmid and a TRAFTAC (either against NFkB or a scrambled control) for 6h, before treating the cells TNF-$\alpha$ (to trigger NFkB activation) and the PROTAC against the Halo tag (HP14) for 19h. The DMSO control sample was the same as a NFkB1, except that it received DMSO instead of the PROTAC.
